# Counting fluorescently labeled proteins in tissues in the spinning disk microscope using single-molecule calibrations

**DOI:** 10.1101/2022.01.02.474734

**Authors:** Maijia Liao, Yin-Wei Kuo, Jonathon Howard

**Affiliations:** Department of Molecular Biophysics and Biochemistry, Yale University, New Haven, Connecticut, USA

**Author notes:** These authors contributed equally to this work (Maijia Liao, Yin-Wei Kuo).

## Abstract

Quantification of molecular numbers and concentrations in living cells is critical for testing models of complex biological phenomena. Counting molecules in cells requires estimation of the fluorescence intensity of single molecules, which is generally limited to imaging near cell surfaces, in isolated cells, or where motions are diffusive. To circumvent this difficulty, we have devised a calibration technique for spinning-disk confocal (SDC) microscopy, commonly used for imaging in tissues, that uses single-step bleaching kinetics to estimate the single-fluorophore intensity. To cross-check our calibrations, we compared the brightness of fluorophores in the SDC microscope to those in the total-internal-reflection (TIRF) and epifluorescence microscopes. We applied this calibration method to quantify the number of EB1-eGFP proteins in the comets of growing microtubule ends and to measure the cytoplasmic concentration of EB1-eGFP in sensory neurons in fly larvae. These measurements allowed us to estimate the dissociation constant of EB1-eGFP from the microtubules as wells as the GTP-tubulin cap size. Our results show the unexplored potential of single-molecule imaging using spinning disk confocal microscopy and provide a straight-forward method to count the absolute number of fluorophores in tissues which can be applied to a wide range of biological systems and imaging techniques.

## Introduction

Measuring the concentration and stoichiometry of macromolecules is essential for quantitatively testing models of dynamic biological processes (Howard, 2014; Pollard, 2014). A potentially general method for counting molecules in living cells is to estimate the intensity of single molecules so that concentrations can be deduced from fluorescence images (Coffman and Wu, 2012; Elf and Barkefors, 2019). This approach has been used successfully to study cell signaling (Mashanov *et al*., 2004; Uyemura *et al*., 2005; Ulbrich and Isacoff, 2007), cytokinesis (Wu and Pollard, 2005), mitosis (Joglekar *et al*., 2006), transcription (Elf *et al*., 2007; Taniguchi *et al*., 2010; Tutucci *et al*., 2018), bacterial motility (Leake *et al*., 2006), and development (Gross *et al*., 2019).

The absolute quantification of single-molecule fluorescence is challenging, however, especially in tissues. Several quantification approaches have been taken, but all have drawbacks. For example, the intensity of fluorescently labeled molecules (such as GFP-tagged proteins) can be calibrated by quantitative immunoblotting (Wu and Pollard, 2005). While this strategy is suitable for cultured cell lines or unicellular organisms, it cannot be applied to tissues due to the difficulty of isolating single cell types for immuno-analysis. A second approach is to visualize single fluorophores, and to use single-step bleaching to estimate the single-molecule fluorescence (Leake *et al*., 2006; Engel *et al*., 2009; Coffman *et al*., 2011). A limitation is that it generally requires a combination of strong illumination and low fluorophore density, for example TIRF imaging near the cell surface (Leake *et al*., 2006; Ulbrich and Isacoff, 2007), and is therefore not suitable for imaging deep in tissues or structures containing larger number of fluorophores. A third approach is fluorescence correlation spectroscopy (FCS), which uses the fluctuations of the fluorescence intensity of the fluorophore-tagged biomolecules in a small volume to obtain quantitative information such as their abundance and mobility (Magde *et al*., 1972; Wachsmuth *et al*., 2015). However, this method is designed for diffusing molecules and is not suitable for molecules undergoing directed motion. Thus, these techniques are either difficult to apply in tissues directly or are restricted to the quantification of biomolecules that are dilute or diffusing.

In this work we visualized single fluorophores in a spinning disk (SDC) microscope, a commonly used instrument for live-cell imaging in tissues, and used single-step photobleaching to calibrate fluorescence intensity. While visualization of single fluorophores has been achieved routinely using total-internal-reflection fluorescence (TIRF), laser-scanning confocal and epifluorescence microscopies, single fluorophore intensities in the SDC microscope have been too low to use them as calibration standards (Lawrimore *et al*., 2011). Here, we compared the fluorescence intensities and photobleaching kinetics in a commercial SDC microscope to those in TIRF and epifluorescence microscopes, to identify the theoretical and practical limitations on the sensitivity of the SDC microscope. The single-fluorophore intensity measured by SDC microscopy serves as a direct calibration standard for fluorophore counting. As a proof of principle, we quantified the number of EB1-eGFP in the EB1 comets and the concentration of free EB1-eGFP in the dendrites of *Drosophila* Class IV dendritic arborization (da) sensory neurons. Our results provide an estimation of the GTP-cap size and the binding affinity of EB1-eGFP, crucial information for characterizing the dynamic and biochemical properties of neuronal microtubules.

## Results and Discussion

### Single fluorophores can be observed by spinning disk confocal microscopy

To quantify molecular fluorescence in a commercial spinning disk confocal (SDC) microscope (Nikon Ti Elipse, Yokagawa spinning disk, 50 μm pinhole, 100 mW diode lasers), we measured the intensities of single fluorophores. Stabilized microtubules (MTs) labeled with a low density of Alexa Fluor 488 were affixed to the surface of the coverslip by anti-tubulin antibodies. We first calibrated molecular fluorescence by total-internalreflection-fluorescence (TIRF) microscopy (Fig. 1A inset), a common method for singlemolecule imaging. The intensities of the fluorescent puncta usually decreased in a single step (Fig. 1A), showing that the majority of puncta corresponded to single Alexa-Fluor-488 dye molecules, as expected. The average intensity of single Alexa Fluor 488 was measured from the histogram of step sizes (Fig. 1B), and the bleaching rate was determined by fitting the fluorescence-lifetime histogram to an exponential (Fig. 1C). Next, we imaged the microtubules by SDC microscopy. We observed fluorescent puncta that resembled the single fluorophores seen by TIRF (Fig. 1D inset). The intensity traces of individual puncta showed clear single-step bleaching events, confirming that the fluorescent signals originated from single fluorophores (Fig. 1D). The fluorophore intensity and bleaching rate were measured from the step-size and lifetime histograms respectively (Figs. 1E and 1F). These results demonstrated that single-molecule imaging by SDC microscopy is possible.

**Figure 1.**
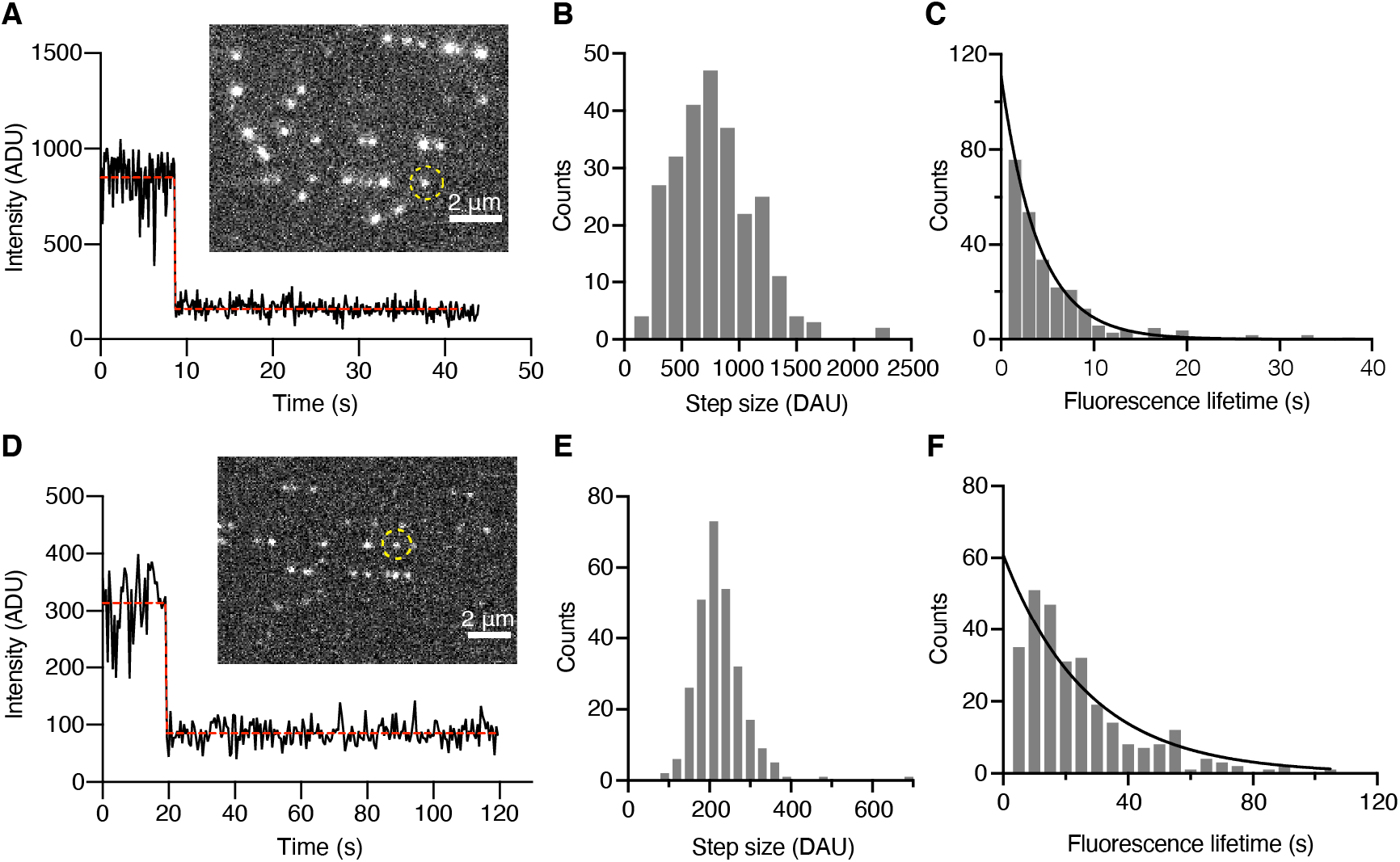
Visualization of single Alexa-Fluor-488 fluorophores by TIRF and SDC microscopies. **A** The black trace shows the intensity as a function of time of a single puncta imaged by TIRF (circled in the inset image). The red dashed line corresponds to a single step identified by the algorithm described in the Materials and Methods. Single-step bleaching can be clearly observed. **B** Histogram of step sizes. **C** Histogram of lifetimes. **D** The black trace shows the intensity as a function of time of single puncta imaged using a Nikon SDC microscope with Yokogawa disk (circled in the inset image). The red dashed line corresponds to a single step. Single-step bleaching can be clearly observed. **E** Histogram of step sizes. **F** Histogram of lifetimes. The sample irradiance was 320 and 290 kW/m^2^ in TIRF and SDC respectively. The images were acquired with 100 ms and 500 ms exposure times in TIRF and SDC respectively.

### Comparison of excitation intensities in the SDC, TIRF and epifluorescence microscopes

The fluorescence intensities of single fluorophores was dimmer in the SDC microscope compared to the TIRF microscope even when the camera exposure time was 5 times longer (under similar excitation laser power, Fig. 1B and 1E). This is due to differences in excitation intensities and emission detection efficiencies. To investigate the excitation intensity differences, we imaged microtubules labeled with a high density of Alexa Fluor 488 in the TIRF, epifluorescence and SDC microscopes. We estimated the intensities of the local excitation fields by measuring the fluorescence bleaching under the same illumination irradiances (the laser power out of the objective divided by the illuminated area). The decay of the fluorescence intensities was well described by single exponentials (see examples in Fig. 2A) and the bleaching rates (obtained from the exponential fit) increased linearly with average illumination irradiance (Fig. 2B), as expected because we are well below the fluorescence saturation intensity (see below).

**Figure 2.**
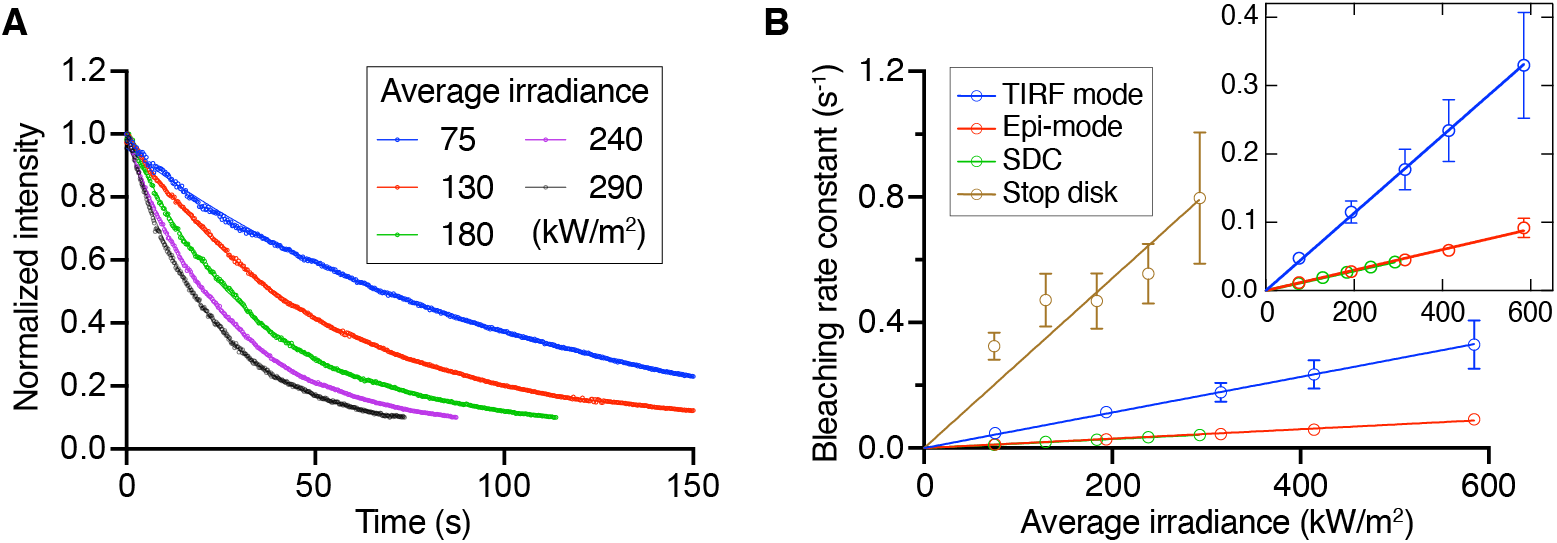
Bleaching rate measurements in different imaging methods. **A** Example bleaching curves under different illumination irradiances in the SDC microscope. The initial segment of each bleaching curve (until ~10% of initial intensity) was fitted to a single-exponential decay (solid lines). **B** Bleaching rate constants derived from the exponential fits increased linearly with irradiance. **Inset**: zoom-in of the bottom three lines. Solid lines: linear regressions constrained to pass through the origin. Error bars: SD from three experimental measurements in each condition.

Under the same illumination irradiances, the bleaching rate constant was 3.8 ± 0.2 (mean ± SE unless otherwise noted) times larger in TIRF compared to epifluorescence microscopy (measured from the slopes of the linear regression, Fig. 2B and Table 2). The roughly four-times higher bleaching rate in TIRF over epifluorescence accords with the ~4-fold enhancement of the evanescent field intensity predicted theoretically (derived in the Supplementary Information and see (Martin-Fernandez *et al*., 2013)). This is the first quantification of enhancement of the evanescence field intensity that we are aware of.

**Table 1.** Slopes of bleaching rate constant vs. irradiance.

| Fit value ± SE | TIRF | Epifluorescence | Spinning disk | Stop disk |
| --- | --- | --- | --- | --- |
| Slope ($10^3 \cdot \text{s}^{-1} \cdot \text{W}^{-1} \cdot \text{m}^2$) | $570 \pm 30$ | $150 \pm 5$ | $144 \pm 2$ | $2700 \pm 200$ |

**Table 2.** Slopes of fluorescence intensity per micron of microtubule vs irradiance.

| Fit value ± SE | TIRF | Epifluorescence | Spinning disk |
| --- | --- | --- | --- |
| Slope<br>(ADU/μm) · W<sup>-1</sup> · m<sup>2</sup>) | 1.91 ± 0.07 | 0.54 ± 0.01 | 0.152 ± 0.003 |

The bleaching rate constants measured from the highly labeled microtubules (TIRF: 5.7 ± 1.0 s; SDC: 24.0 ± 1.1 s; Table 3, details in Materials and Methods) were in good agreement with the bleaching rate constants obtained from the single-molecule lifetimes (TIRF: 5.3 ± 1.1 s; SDC: 24 ± 3 s). This is further confirmation of the reliability of our single-molecule SDC observations.

Under the same illumination irradiances, the bleaching rate constants of SDC and epifluorescence were similar (ratio of slopes = 0.96 ± 0.04) (Figure 2B, Table 2). This shows that the excitation field intensities are similar in the two imaging modes when the irradiance is the same, as expected.

In SDC, each point in the sample is illuminated intermittently due to the rotation of the spinning disk; therefore, only a small percentage of the total area is illuminated at any one time. To assess the effect of the disk rotation, we measured the bleaching rate in illuminated regions when the disk was stationary (i.e., the stop-disk condition). When the disk was stationary, the bleaching rate was 19 ± 1 times larger than in the spinning-disk mode (Fig. 2B and Table 1). The instantaneous irradiance of each spot was thus about twenty times larger than the average irradiance, and only ~5% of the field of view was illuminated by the excitation light when the disk was spinning. This result confirms that most of the illumination light is blocked by the spinning disk, transmitting only a small portion of excitation light at each point of the sample.

In summary, the higher fluorescence intensities in the TIRF microscope are due in part to the 4-fold enhancement of the intensity of the evanescent filed that excites the fluorophores. The excitation intensities in epifluorescence and SDC are similar when the instantaneous illumination in the SDC is increased roughly 20-fold to compensate for the attenuation due to the small fraction of the spinning disk occupied by the pinholes.

### Comparison of emission efficiencies in the SDC, TIRF and epifluorescence microscopes

We next compared the fluorescence intensities of the highly labeled Alexa-Fluor-488 microtubules when imaged by the TIRF, epifluorescence and SDC microscopes. Consistent with the single-molecule measurements, fluorescent microtubules were the brightest by TIRF microscopy and dimmest by SDC microscopy (Fig. 3A) under similar illumination irradiances and identical camera exposure times. Like the bleaching rates, the fluorescence intensities increased linearly with increasing illumination irradiance (Fig. 3B), showing that the excitation light intensity is within the linear dynamic range. This is expected since the excitation intensity we used was much smaller than the predicted saturation intensity of Alexa Fluor 488 (4 GW/m^2^ – this is about 10000-times higher than the maximum irradiance we used). From the slopes of the linear regression, TIRF microscopy is 3.5 ± 0.2 times brighter than epifluorescence (Table 2), which agrees well with the 3.8 ± 0.2 fold increase in TIRF’s excitation field measured from the bleaching rate (Table 1), and is expected from the enhancement of the evanescent field intensity in TIRF (mentioned above). However, the SDC fluorescence intensity was 3.6 ± 0.1 fold lower than epifluorescence (Fig. 3B and Table 2), even though the excitation intensities were similar (as judged by the similar bleaching rates stated above). The lower intensities of the SDC images compared to epifluorescence indicates that the SDC microscope suffers more light loss in the emission pathway. The loss is not due to the objective as the same objective was used. Some of the loss is expected to be due to the pinhole (~20%, Supplementary Fig. S1, (Wilson, 1990)); the rest of the loss is presumably due to the lenses, mirrors, and filters. In summary, we have accounted for the lower fluorescence intensity measured in SDC microscopy compared to in widefield (epifluorescence and TIRF) microscopy as a combination of different excitation intensities and emission efficiencies.

**Figure 3.**
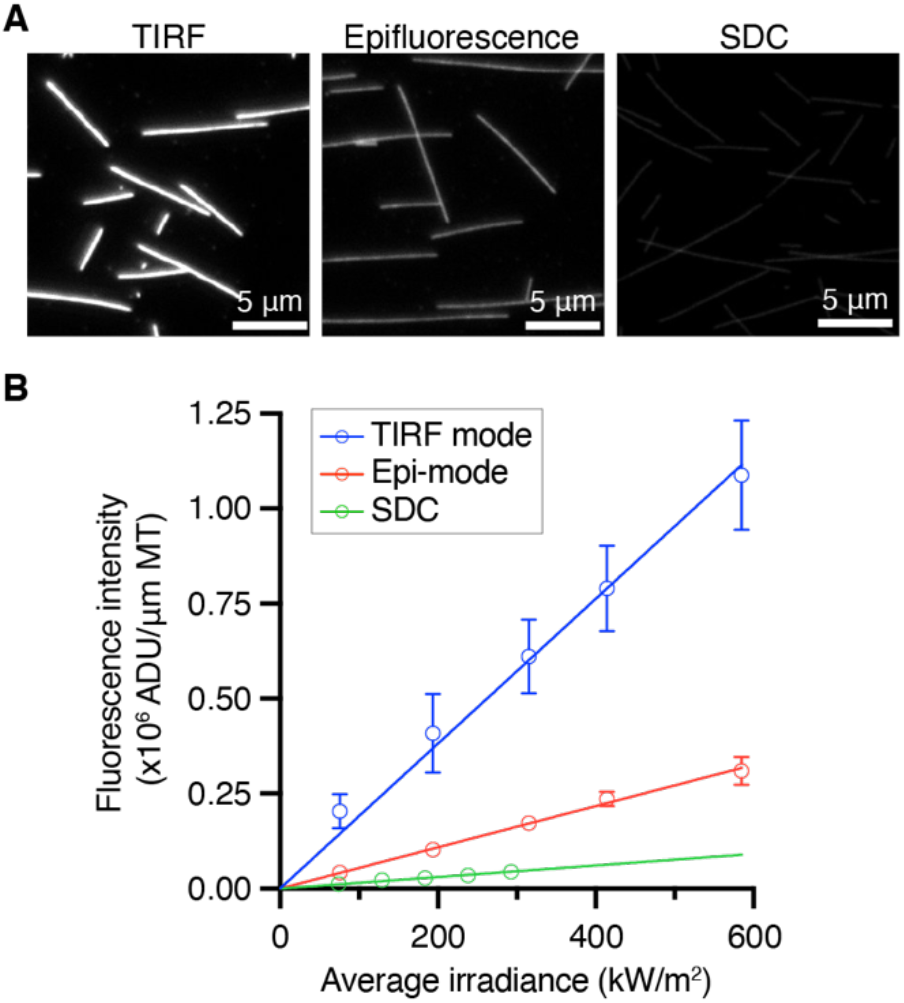
Fluorescence intensities of Alexa-Fluor-488 microtubules. **A** Example images of Alexa-Fluor-488 microtubules imaged by TIRF, epifluorescence and SDC microscopy under similar average illumination irradiances (76 kW/m^2^ for TIRF and epifluorescence; 75 kW/m^2^ for SDC) with the same camera exposure time (100 ms). Images were adjusted to the same contrast level. **B** Fluorescence intensities increased linearly with average irradiance in all three imaging methods. Error bars: SD. Each condition was measured from 3 experimental measurements. Solid lines: linear regression of each imaging method. The linear regression is constrained at the origin.

As an overall check on our single-molecule fluorescence calibrations, we estimated the density of the Alexa Fluor 488 in the highly labeled microtubules. The density was 59% (estimated by TIRF) and 54% (estimated by SDC); these values are similar to the labeling density estimated by absorbance using a spectrophotometer (43 ± 1%; mean ± SD) (Table 3). Therefore, we have established a calibration standard for fluorophore counting by measuring the bleaching step size of single fluorophores using SDC microscopy.

**Table 3.** Single-fluorophore fluorescence intensity and lifetime in TIRF and SDC microscopy.

| Mean ± SD* | TIRF single fluorophore | TIRF high fluorophore density MT | SDC single fluorophore | SDC high fluorophore density MT | Tetraspeck bead (TIRF) | Tetraspeck bead (SDC) |
| --- | --- | --- | --- | --- | --- | --- |
| Bleaching time (s) | 5.3 ± 1.1 | 5.7 ± 1.0 | 24 ± 3 | 24.0 ± 1.1 | n/a | n/a |
| Step size per 100 ms exposure time (ADU) | 600 ± 160 | - | 47 ± 3 | - | - | - |
| Intensity per μm of MT ADU for 100 ms exposure time | - | 6.1 ± 1.0 $\times 10^5$ | - | 4.4 ± 0.2 $\times 10^4$ | - | - |
| Total Intensity per bead per 100 ms exposure time (ADU) | - | - | - | - | 1.34 ± 0.23 $\times 10^5$ | 1.06 ± 0.04 $\times 10^4$ |
| Average irradiance (W/cm<sup>2</sup>) | 32 | 32 | 29 | 29 | 32 | 29 |
| Estimated fluorophore labeling density** | 59% |  | 54% |  | n/a |  |

\*SD from n = 3 measurements
\*\*Fluorophore density estimated by absorbance is $43 \pm 1\%$ ( $n = 3$ measurements).

### Establishing a consistent calibration standard for fluorophore counting with SDC microscopy

To facilitate calibration of the single-molecule fluorescence in the SDC microscope, we used fluorescent beads as a reference standard. We imaged TetraSpeck beads (Thermofisher, cat no.T7279) in both the TIRF and SDC microscopes under the same imaging conditions as the single-fluorophore experiments. The intensity ratios of TetraSpeck beads to single Alexa-Fluor-488 dyes measured by TIRF and SDC were 225 ± 72 (mean ± SE) and 227 ± 18 respectively (Table 3). By periodically measuring the intensity of TetraSpeck, we could test whether there has been a change in the excitation or emission intensities of the SDC microscope.

When using different fluorophores, we adjusted the calibration based on the relative brightness compared to Alexa Fluor 488 at the illumination wavelengths, as well as considering the sample irradiance (laser intensity setting) and the camera exposure time.

The accuracy of our calibration is primarily determined by the accuracy of the step size measured in the SDC, which had a coefficient of variance of 0.07 (SD/mean in three independent measurements). From the *t*-distribution, the 95% confidence interval is ±0.07 × 2.92 ≅ 0.2 (2 degrees of freedom). Other sources of error include measurement of the irradiance (< 5% SD), and uncertainties of the fluorescence intensities inside the cell due to molecular interactions. These latter effects are difficult to judge. Overall, we expect our calibration to be accurate to better than 20% (SD/mean), which make the 95% confidence range roughly a factor of two (between 0.6 and 1.4).

### Quantification of EB1-eGFP in *Drosophila* neurons

Microtubule end-binding proteins (EBs) track growing microtubule ends *in vitro* and *in vivo*. The amount and the size of the EB binding zone on a growing microtubule tip can serve as an indicator of the GTP-tubulin cap size (Zanic *et al*., 2009; Maurer *et al*., 2011; Seetapun *et al*., 2012; Coombes *et al*., 2013; Strothman *et al*., 2019; Roostalu *et al*., 2020). Additionally, EB proteins can recruit various microtubule-associated proteins that may be critical for the dynamics and function of microtubule filaments (Akhmanova and Steinmetz, 2015). While the number and direction of EB comets are frequently used to estimate the number and polarity of growing microtubules in neurons (Stone *et al*., 2008; Stewart *et al*., 2012; del Castillo *et al*., 2015), the absolute number and binding region of EB molecules have not been examined in detail. Quantifying the amount of EB proteins at a growing microtubule end, the size of the EB-binding region (i.e., the EB-cap size) and its binding affinity thus provide valuable insights into the dynamical properties of the microtubule cytoskeleton.

To quantify EB comets *in vivo*, we expressed EB1-eGFP and the cell membrane marker CD4-tdTomato in class IV neuron of *Drosophila* larvae under the cell-specific promoters (Rolls *et al*., 2007; Han *et al*., 2011). EB1 preferentially binds to growing microtubule ends and turns over rapidly; in this way it forms a comet-like distribution behind the polymerizing tip of the microtubule (Bieling *et al*., 2007; Dixit *et al*., 2009). Imaged by SDC microscopy, fluorescent comets were observed throughout the dendrites of these neurons (example in Fig. 4A). To measure the shapes of the comets, we modeled the comets as exponential decays convolved with a Gaussian point-spread function using a Monte-Carlo optimization procedure (red dashed curve in Figure 4B and see Fig. S3 to S5 and the Methods section for details). By considering the fluorophore brightness (i.e., the product of extinction coefficient and quantum yield), emission spectra, illumination irradiance and camera exposure time, we determined the intensity of a single eGFP using the bleaching step size of Alexa Fluor 488 as a calibration standard (see Methods for details). We found that each comet contains on average 84 ± 25 (mean ± SD, *n* = 6 larvae) EB1-eGFP (Fig. 4C), with an exponential decay length *λ* of 190 ± 40 nm. If we assume that EB1 binds to GTP-tubulin in the lattice, then the GTP cap has an average length of 190 nm (where the density decreases e-fold). Given that there are 300 tubulin dimers in 190 nm (assuming the dimer length is 8.2 nm and there are 13 protofilaments and that the GTP-tubulin density decays exponentially from 100% at the end), we estimate that 28% of the GTP-tubulins have EB1-eGFP bound (84/300).

**Figure 4.**
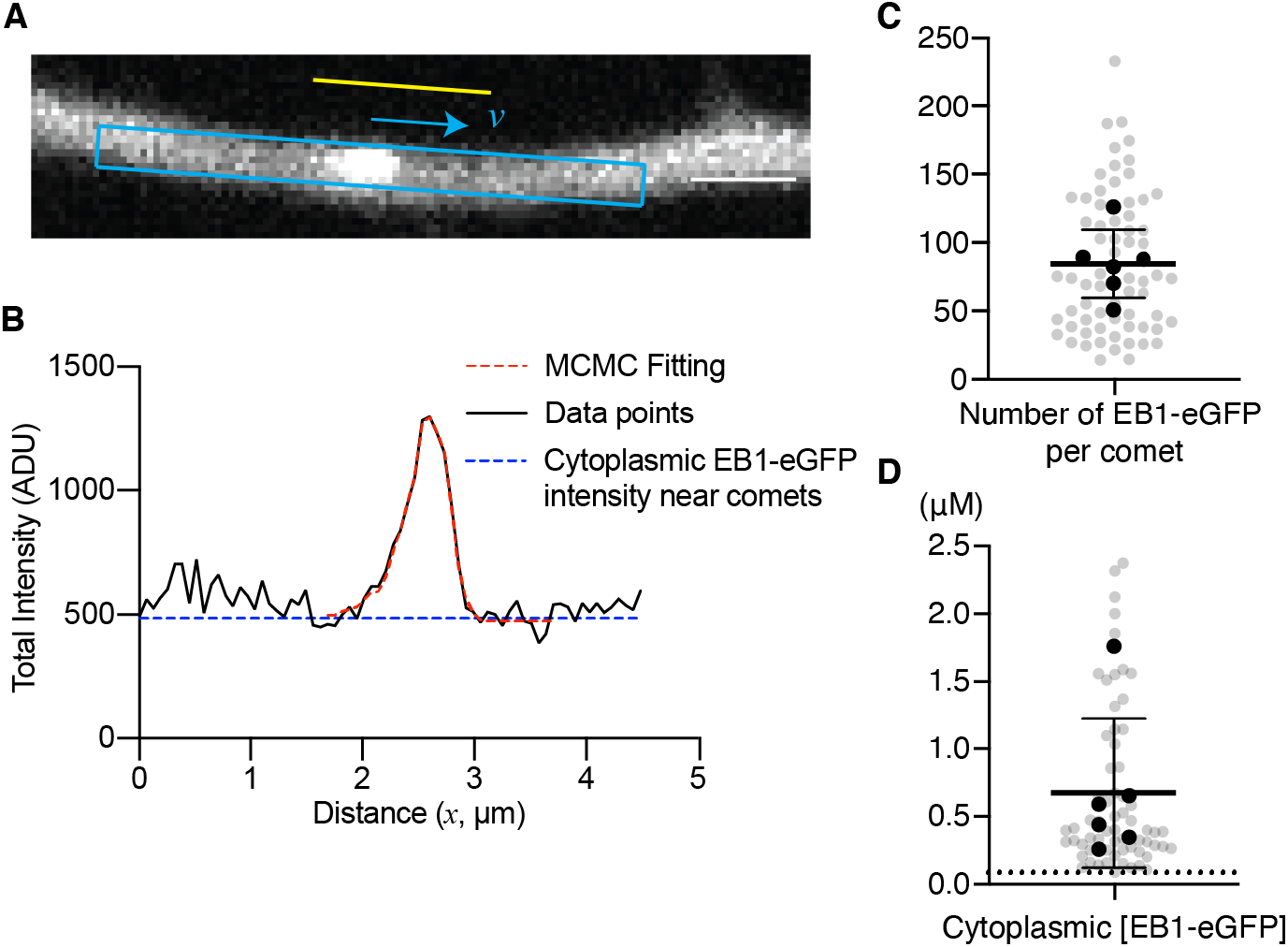
Determination of the amount of EB1-eGFP at microtubule ends and cytoplasm. **A** An example of an EB1-eGFP comet. The overall intensity was calculated within the 0.5 μm × 4.5 μm blue rectangle. The intensity of adjacent regions outside of the dendrite (yellow line) was used for background subtraction. Scalebar 1 μm. **B** The measured intensity profile (black) was modeled as an exponential convolved with a Gaussian point-spread function (red dashed line, see Materials and Methods). The offset (blue line) is considered to be the cytoplasmic EB1-eGFP intensity. **C** Number of EB1-eGFPs per comet. **D** Concentration of cytoplasmic EB1-eGFP. Black points indicate the average from each larvae (6 larvae; data points from all 68 comets were shown in light gray). The minimum concentration that can be detected (signal-to-noise ratio > 3) was ~ 0.09 μM (dashed line in D). Mean and SD from 6 larvae are shown as horizontal bars.

The cytoplasmic EB1-eGFP concentration was 0.68 ± 0.55 μM (Fig. 4D; mean ± SD, n = 6 larvae; see methods for details), which is within the large range of physiological EB1 concentration reported from various species and cell types (from ~0.14 μM in budding yeast to 2.1 μM in HeLa cells)(see references in Supplementary Table S1). Combining these measurements, we can estimate the binding affinity between EB1-eGFP and GTP-tubulin lattice by: 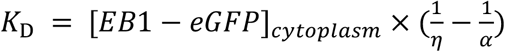 (assuming that unlabeled EB1 and EB1-eGFP have the same affinities), where *K_D_* is the dissociation constant, *η* is the occupancy of EB1-eGFP (the number of EB1-eGFP bound divided by the total GTP-tubulin sites in the lattice) and *α* is the fraction of EB1 containing eGFP (i.e. the labeling density). If we assume *α* is equal to 1, *K_D_*=1.6 ± 0.7 μM (mean ± SD, *n* = 6 larvae), which is similar to the dissociation constant estimated in human tissue culture cells (3.8 μM, (Seetapun *et al*., 2012)), but much larger than the one reported from *in vitro* measurements (22 nM, (Maurer *et al*., 2014)) (Supplementary Table S2). This estimation represents an upper bound of *K_D_* since we assumed that the overexpressed EB1-eGFP is much more abundant than the endogenous unlabeled EB1. Thus, our results suggest that these neurons expressed a few hundred nM of EB1-eGFP, with micromolar-level binding affinity to the GTP-tubulin lattice in dendrites.

Previous *in vitro* studies have shown that the decay length of EB comets increases with the microtubule polymerization rate (Bieling *et al*., 2007; Strothman *et al*., 2019; Farmer *et al*., 2021). To investigate whether this correlation holds for dendritic microtubules, we examined the relationship between the number of EB1-eGFP, the decay length of EB1-eGFP comet and the comet speed. The comet velocity (0.094 ± 0.057 μm/s, mean ± SD from 68 comets in 6 larvae, similar to previously reported EB comets speed in *Drosophila* neurons (Ori-McKenney *et al*., 2012; Poe *et al*., 2017)) showed positive correlations with both decay length and EB1-eGFP number (Pearson’s *r* = 0.53 and 0.55 respectively, *n* = 68 comets), suggesting that faster growing microtubules have larger GTP-caps and a larger number of bound EB1-eGFP (Fig. 5A, B). These correlations are consistent with previous measurements in a cultured human cell line (Matov *et al*., 2010) and in the axons of cultured primary neurons from *Drosophila* (Hahn *et al*., 2021). Our measurements provide additional information, namely the absolute number and affinity of EB1-eGFPs in tissues rather than cultured cells.

**Figure 5.**
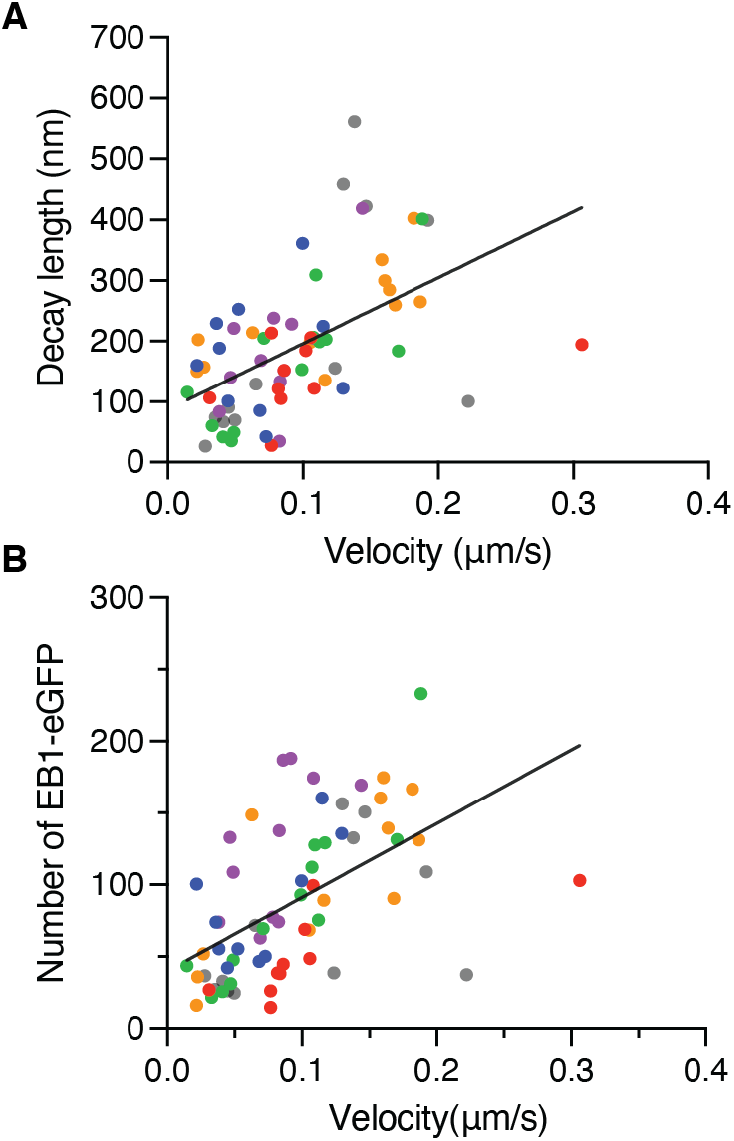
Comet velocity positively correlates with the comet decay length. **A** The comet length correlates with velocity. **B** The number of EB1-eGFPs correlates with velocity. Pearson correlation coefficients are 0.53 and 0.55 respectively (two-tailed Pearson correlation test, *p* < 0.0001, *n* = 68 comets). Comets from 6 larvae were shown in different colors.

The GTP-cap size of the dendritic microtubules estimated here (~23 layers of tubulin) was only about half of the GTP-cap size estimated in the cultured human LLCPK1 epithelial cells (~55 layers) (Seetapun *et al*., 2012), potentially reflecting the slower average speed of the dendritic EB comets (94 nm/s in the dendrites, 157 nm/s in LLCPK1 cells; mean ± SE, see Supplementary Table S2 for summary). Interestingly, the EB-comet lengths observed in the dendrites of *Drosophila* neurons (188 ± 14 nm with a polymerization rate of 94 ± 7 nm/s; mean ± SE) was considerably smaller than the comet lengths of reconstituted microtubules with similar growth rate polymerized from purified mammalian brain tubulin (Maurer *et al*., 2014; Chaaban *et al*., 2018; Strothman *et al*., 2019) (summarized in Supplementary Table S2). Instead, the comet length is comparable to those on microtubules polymerized from *C. elegans* tubulin; these microtubules displayed higher catastrophe frequency than those from mammalian brain tubulin (Chaaban *et al*., 2018). While the GTP-cap size may not be the sole factor to determine the catastrophe frequency (Bowne-Anderson *et al*., 2015; Farmer *et al*., 2021), the shorter cap size may imply more frequent shrinkage events in these dendritic microtubules. Live imaging of single microtubule filaments in the neuron of a whole organism remains a major challenge, leaving the direct measurement of neuronal microtubule dynamics difficult. Future investigation of the EB cap size in mutants of microtubule polymerases, depolymerases and microtubule stabilizers can potentially provide new insights into the relationship between GTP-cap size and microtubule stability in these neurons.

## Conclusion

We introduced a strategy to quantify the fluorophore number in tissue using spinning-disk-confocal microscopy. We imaged single fluorophores in the SDC microscope and used the single-fluorophore bleaching step size to calibrate fluorophore numbers. Applying this strategy, we quantified the number of EB1-eGFP in both puncta-like structures (i.e., EB1 comets) as well as its cytoplasmic concentration, allowing the estimation of binding affinity *in vivo*. This approach is built on the assumption that the fluorophore intensities inside the cells are similar to the ones measured *in vitro*, while the brightness of fluorophores may depend on the environments. More detailed intensity corrections can be performed based on previous reports comparing the photoproperties of several fluorescent proteins from both *in vitro* and *in vivo* systems (Chen *et al*., 2002; Heppert *et al*., 2016; Botman *et al*., 2019) to achieve better accuracy. A major advantage of the strategy introduced here is that the calibration process does not require additional microscopic or biochemical methods and can be performed directly using the identical microscopy setup as the *in vivo* imaging experiments. The calibration procedure can be simplified without the needs of extra imaging standards (e.g., fluorescent beads or purified protein standards used for immunofluorescence), tissue fixation or the isolation of specific cell types from the tissues. Additionally, our fluorescent intensity and bleaching rate measurements using TIRF, epifluorescence and SDC microscopy provide a quantitative assessment of the minimal brightness required for visualizing single-fluorophore using SDC microscopy, which is an important step toward single-molecule imaging in cellular systems. The quantitative imaging methods introduced here is broadly applicable for quantifying the number of target molecules in the live cells from tissues, which are typically more challenging systems for other fluorophore counting methods.

## Materials and Methods

### Flow chamber and microtubule preparation

Tubulin was purified from bovine brain as previously described (Castoldi and Popov, 2003). The preparation of imaging chambers and stabilized microtubules followed the methods described in (Gell *et al*., 2010; Kuo and Howard, 2021). All reagents were purchased from Sigma-Millipore except otherwise noted. To affix the microtubules onto the surface of the silanized coverslips, 25 μg/mL of anti-tubulin antibody (clone SAP.4G5) solution was perfused in the flow chamber with 5 minutes incubation, and washed by BRB80 buffer (80 mM PIPES-KOH, pH 6.9, 1 mM EGTA, 1 mM MgCl_2_). The channel was then passivated by incubating with 1% pluronic F127 solution, followed by 2 mg/mL casein solution as previously described (Kuo *et al*., 2019). Alexa-Fluor-488-labeled stabilized microtubules was prepared by polymerizing Alexa-Fluor-488-conjugated bovine tubulin in the presence of slowly hydrolysable GTP analog GMPCPP (Jena Bioscience) as previously described (Gell *et al*., 2010). For single-fluorophore imaging, Alexa-Fluor-488 tubulin was mixed with unlabeled tubulin so that the final labeling density of the fluorophore was around 0.09%. Oxygen scavenger solution (40 mM glucose, 40 μg/mL glucose oxidase, 16 μg/mL catalase, 0.1 mg/mL casein, 1% β-mercaptoethanol in BRB80) was used for all fluorescent microtubule imaging experiments.

### Microscopy setup

TIRF and epifluorescence imaging was performed on an inverted microscope (Nikon Ti Eclipse), with a 488 nm excitation laser and a 525/50 nm emission filter. For spinning disk confocal imaging, an inverted Nikon TI microscope equipped with a confocal scanner unit (CSU-W1 disk, Yokogawa) which contains a four-bands dichroic beamsplitter (Di01-T405/488/568/647, Semrock) and a 525/50 nm emission filter. Both microscopes were operated by Nikon NIS element software. All images were collected by a 100x/1.45 NA oil objective (CFI Plan Apochromat Lambda, Nikon) with sCMOS cameras (Zyla 4.2 plus, Andor).

### Fly stocks

We used Gal4 driver line *ppk-Gal4* to drive the expression of UAS-EB1-eGFP (Stock 35512 from Bloomington *Drosophila* Stock Center) to visualize the growing plus ends of microtubules and the reporter line, *ppk-CD4-tdTomato* (Han *et al*., 2011) to observe the dendrite morphology of Class IV da neurons.

### Larvae sample imaging

Embryos were collected for 2 hours on apple juice agar plates with a dollop of yeast paste and aged at 25 °C in a moist chamber. The plates containing the first batch of embryos were discarded as the dendrite morphology of da neurons is less consistent in those animals. Larvae were immobilized individually on agarose pads (thickness 0.3-0.5mm) sandwiched between a slide and a coverslip. The imaging was done using a spinning disk microscope: the Yokogawa CSU-W1 disk (pinhole size 50 μm) built on a fully automated Nikon TI inverted microscope with perfect focus system, an sCMOS camera (Zyla 4.2 plus sCMOS), and running Nikon Elements software.

### Measurements of fluorescence intensity and bleaching kinetics

The pixel intensities within 7×7 pixel^2^ boxes (450×450 nm^2^) centered at each tetraspeck bead or single fluorophore were summed to get the overall intensity. The background signal was estimated from regions away from fluorescent spots and further subtracted from the overall intensities.

Fluorescent intensity traces for individual Alexa-Fluor-488 dyes were obtained by calculating overall intensities over time. To analyze the single fluorophore photobleaching data, we used the previously developed step detection algorithm that use statistical tests based on the two-sample *t* test without assumed equal variance to identify steps (Chen *et al*., 2014). The detected steps smaller than a quarter of the average single bleaching step size were merged with neighboring steps to avoid unrealistic small steps. Steps that last longer than 6 frames and with signal to noise ratio larger than 3 were used for step size and single step bleaching time estimations.

### Single fluorophore intensity estimation

The intensity ratio of single eGFP and Alexa Fluor 488 (*l_eGFP_/l_Alexa488_)* on SDC microscopy can be estimated based on the following equation:

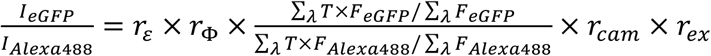, where *r_ε_* indicates the ratio of extinction coefficients at excitation wavelength (488 nm); r_Φ_ is the ratio of quantum yields; *T* is the overall transmittance of the emission filter and dichroic beamsplitter; *F* is the relative emission intensity of the two fluorophores; *r_cαm_* and *r_ex_* corresponds to the ratios of camera exposure time and excitation irradiance between the imaging conditions from the *in vivo* experiments and *in vitro* single-molecule calibration in SDC microscope respectively. The extinction coefficients and quantum yields were obtained from Thermofisher and the Fluorescent Protein Database (FPbase). The emission spectra were obtained from Chroma Spectral Viewer and the transmittance profiles of the dichroic mirror and emission filters were based on the specifications from manufacturers. The conversion factor from the spectral properties of fluorophores in our system=0.65.

### Cytoplasmic EB1-eGFP concentration estimation

In axial dimension, diffraction limits the resolution to *2nλ/NA*^2^ with *n* the refractive index of the medium between objective and sample, corresponding to a depth of field~ 800nm. The thickness of most branches studied in this paper fall within the estimated depth of field. Thus, we used the plateau of the MC fitting of the fluorescence signal (Fig. 4B, blue dashed line) from the mid-plane of the dendrite to calculate the cytoplasmic EB1-eGFP concentration. The cytoplasmic EB1-eGFP concentration can be obtained by dividing the total number of local cytoplasmic EB1-eGFP by the volume of the dendrites approximated by 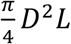 (where D is the FWHM of the dendrite from the membrane marker and *L* corresponds to the length of the dendrite where total intensity was extracted).

### MC optimization method

The overall intensity profile of the EB1-eGFP comets within 0.5×4-8 μm^2^ rectangular box can be fitted by 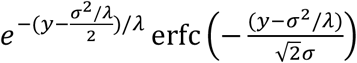 obtained through convolution of gaussian with exponential decay, where *σ* stands for the standard deviation of gaussian and *λ* is the exponential decay length. The fitted parameters are used to set the range of *σ* and *λ* which are used as inputs for MC optimizations (code details in Github: https://github.com/Maijia-cpu/Comet-profile). In each step of MC optimization method, a simulated image is generated based on 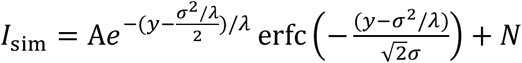, where *A* stands for the image intensity, *N* stands for image photon shot noise. The MC optimization aim to find the *σ* and *λ* that minimize the square of the difference (Liao *et al*., 2021): 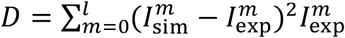, where *I*_sim_ and *I*_exp_ stand for 1D intensity profiles for simulated and experimental images separately, *m* stands for pixel number. Six 1000-step simulations were carried out for *σ* and *λ* detection. The obtained *σ* and *λ* are averages of results from six simulations. The total intensity of the comet was calculated by summing the total experimentally measured signal within the range of −2σ to 2*λ* with zero being the center of the comets.

## Author contributions

M.L and Y.-W.K. performed all experiments and data analysis, all authors designed research and wrote the paper.

## Acknowledgement

We thank Dr. Mohammed Mahamdeh for the discussions on the work and the Howard lab members for the feedback on the manuscript. This work was supported by NIH Grants R01 NS118884 and R01 GM139337 (to J.H.).

