## Supplementary Information for "Counting fluorescently labeled proteins in tissues in the spinning disk microscope using single-molecule calibrations"

**Table S1.** Physiological concentration of EB proteins in different organisms and cell types

| Organism/cell types | Concentration | Reference |
|---|---|---|
| *Saccharomyces cerevisiae* (Bim1 in budding yeast) | ~140 nM (copy number of Bim1 from Ghaemmaghami et al.; median cell volume of 42 fL based on Jorgensen et al.) | (Jorgensen *et al.*, 2002; Ghaemmaghami *et al.*, 2003) |
| Xenopus egg | 0.24 µM | (Tirnauer *et al.*, 2002) |
| *Schizosaccharomyces pombe* (Mal3 in fission yeast) | ~200 nM | (Alberico *et al.*, 2013) |
| HeLa cells | 2.1 µM | (Itzhak *et al.*, 2016) |

**Table S2.** Summary of EB cap size, MT polymerization rates and dissociation constant

| EB cap size (decay length; mean±SE) | MT growth rate (mean±SE) | Systems | EB affinity ($K_D$) | Reference |
|---|---|---|---|---|
| 188±14 nm | 94±6.9 nm/s | Dendrites of *Drosophila* class IV *da* neuron | 2.25±0.47 µM | This study |
| 440±26 nm | 156.6 ±13 nm/s | LLCPK1 epithelial cells | 3.8 µM | (Seetapun *et al.*, 2012) |
| 383±10 nm | ~82 nm/s | *In vitro* reconstitution (Human EB1, bovine tubulin) | N/A | (Strothman *et al.*, 2019) |
| 276±14 nm | 35±3 nm/s | | | |
| ~ 230 nm | 49 nm/s | *In vitro* reconstitution (human EB1, porcine tubulin) | 22±1 nM | (Maurer *et al.*, 2014) (Decay length estimated from Fig. 2F) |
| ~ 330 nm | 71 nm/s | | | |
| ~ 380 nm | 95 nm/s | | | |
| 339±22 nm | 87±5 nm/s | *In vitro* reconstitution (human EB1, bovine tubulin) | N/A | (Chaaban *et al.*, 2018) |
| 144±20 nm | 88±5 nm/s | *In vitro* reconstitution | N/A | |

| | | | | |
|---|---|---|---|---|
| | | (human EB1, C. elegans tubulin) | | |
| 610±20 nm | ~170 nm/s | *In vitro*, human EB1, bovine MT | N/A | (Farmer *et al.*, 2021) |
| 650±20 nm | | *In vitro*, human EB1, bovine MT with XMAP215 | | |

**Evanescent Electric Field for Total Internal Reflection**

Based on the boundary conditions that normal components of $\boldsymbol{D}$ and $\boldsymbol{B}$ are continuous and tangential components of $\boldsymbol{E}$ and $\boldsymbol{H}$ are continuous.

The reflected and transmitted amplitudes are related to the incident amplitude through Fresnel equations. The ratios *r* and *t* are the Fresnel reflection and transmission amplitude coefficients respectively defined as:

$$r = \frac{E_r}{E_{in}}$$

$$t = \frac{E_t}{E_{in}}$$

for *s*-polarized incident light:

$$r^s = \frac{ncos\theta_i - \sqrt{n'^2 - n^2 sin^2\theta_i}}{ncos\theta_i + \sqrt{n'^2 - n^2 sin^2\theta_i}}$$

$$t^s = \frac{2ncos\theta_i}{ncos\theta_i + \sqrt{n'^2 - n^2 sin^2\theta_i}}$$

and similarly for *p*-polarized incident light,

$$r^p = \frac{n'^2 cos\theta_i - n\sqrt{n'^2 - n^2 sin^2\theta_i}}{n'^2 cos\theta_i + n\sqrt{n'^2 - n^2 sin^2\theta_i}}$$

$$t^p = \frac{2nn' cos\theta_i}{n'^2 cos\theta_i + n\sqrt{n'^2 - n^2 sin^2\theta_i}}$$

To fully describe the evanescent field at interface, we write the transmitted electric field in vector form $\boldsymbol{E}_{tp} = (E_{tpx}, E_{tpy}, E_{tpz})$ at the boundary.

$$E_{tpy} = E_{in}^{s} t^{s}$$

$$E_{tpx} = E_{in}^{p} t^{p} cos\theta_t$$

$$E_{tpz} = E_{in}^{p} t^{p} sin\theta_t$$

$$n' sin\theta_t = n sin\theta_i$$

For totally internal reflected light, $sin\theta_i > sin\theta_c = {n'}/{n}$.

$$t^p = \frac{2nn' cos\theta_i e^{-i\delta^p}}{\left(n'^4 cos^2\theta_i + n^4 sin^2\theta_i - n^2 n'^2\right)^{1/2}}$$

$$t^s = \frac{2n cos\theta_i e^{-i\delta^s}}{\left(n^2 - n'^2\right)^{1/2}}$$

$$\delta^p = arctan\left[\frac{n\sqrt{n^2 sin^2\theta_i - n'^2}}{n'^2 cos\theta_i}\right]$$

$$\delta^s = arctan\left[\frac{\sqrt{n^2 sin^2\theta_i - n'^2}}{n cos\theta_i}\right]$$

$$E_{tpy} = E_{in} \frac{2n cos\theta_i}{\left(n^2 - n'^2\right)^{1/2}} e^{-i\delta^s}$$

$$E_{tpx} = E_{in} \frac{2n cos\theta_i (n^2 sin^2\theta_i - n'^2)^{1/2}}{\left(n'^4 cos^2\theta_i + n^4 sin^2\theta_i - n^2 n'^2\right)^{1/2}} e^{-i(\delta^p + \pi/2)}$$

$$E_{tpz} = E_{in} \frac{2n^2 cos\theta_i sin\theta_i}{\left(n'^4 cos^2\theta_i + n^4 sin^2\theta_i - n^2 n'^2\right)^{1/2}} e^{-i\delta^p}$$

The evanescent field intensity at the dielectric interface as a function of the incident field is:

$$I_t^s = {E_{tpy}}^2 = I_i^s \frac{4n^2 cos^2\theta_i}{n^2 - n'^2}$$

$$I_t^p = {E_{tpx}}^2 + {E_{tpz}}^2 = I_i^p \frac{4n^2cos^2\theta_i\left(2n^2sin^2\theta_i - n'^2\right)}{n'^4cos^2\theta_i + n^4sin^2\theta_i - n^2n'^2}$$

where $I_i^s = {E_{in}^s}^2$ and $I_i^p = {E_{in}^p}^2$.

For glass-water interface we studied, $n' = 1.33$ and $n = 1.515$. $sin\theta_i = 1.33/1.515$. Thus we have:

$$I_t^s = 4I_i^s$$

$$I_t^p = 5.19I_i^s$$

The evanescent field intensity is greater than the incident intensity, thus improving the signal to noise ratio.

### Estimation of pinhole effect

The equation defining PSF of a standard confocal microscope is:

$$PSF_{conf}(x,y,z) = PSF_{exc}(x,y,z) \cdot \left(PSF_{det}(x,y,z) \otimes D(x,y)\right) \quad (S1)$$

where $PSF_{exc}$ and $PSF_{det}$ are point spread functions of excitation and detection respectively. $D$ is a function modeling pinhole shape (here we use circular aperture). The convolution operator acts only on the lateral coordinate $(x,y)$. The confocal point spread function $PSF_{conf}$ is given by the product of excitation and collection PSFs, with the latter degraded by convolution of the pinhole. Fig. S1 shows the total intensity at focal plane as a function of pinhole diameter.

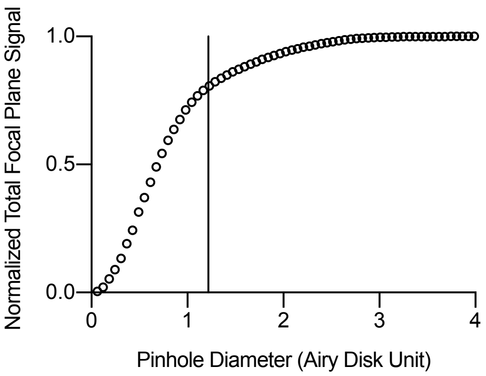


**Fig. S1** The dependence of the total focal plane signal with pinhole diameter. $PSF_{exc}$ and $PSF_{det}$ are modeled by Gibson and Lanni model.

The first minimum of the Airy disk from a point source is: $1\ Airy\ disk\ unit = 1.22\frac{M_{obj}\lambda_{ex}}{NA}$. The pinhole diameter we used in experiment is 50μm (1.22 Airy disk unit shown by the black line in Fig. S1). The total intensity obtained with 50 μm pinhole reaches 80% of the total amount of light (i.e., pinhole size goes to infinity).

### The crosstalk factor is negligible

The larvae with genotype: ; *ppkgal4/+;ppkCD4tdTomato/UAS-EB1-eGFP* were used for visualizing EB1 comets as well as dendrites in ddaC neurons. We focused on third-instar larvae

96 hr after egg laying and observe the dynamics of comets using spinning disk confocal microscope.

Bleed-through can be a potential issue when simultaneously imaging EB1-eGFP (green channel) and CD4tdTomato (red channel). To estimate the bleed-through from red to green channel when calculating EB1-EGFP intensities, we imaged the larvae with CD4tdTomato expressed alone (genotype: *;ppkgal4; ppkCD4tdTomato*) in both red and green channels and built a cross-talk calibration curve. A polynomial function was used as a model to establish the relationship between the bleed-through intensity in the green channel and the intensity in the red channel. The coefficient of the polynomial and the red channel intensity were used to subtract the bleed-through contribution in the green channel.

Figure S2 shows the intensity in the red channel and the corresponding bleed-though intensity in the green channel for different positions in CD4tdTomato expressed dendrites. Coefficients from the polynomial fitting model were used to predict and subtract the bleed-through contribution in the red channel. The red channel pixel intensity ranges from 50-250 ADU. Based on the polynomial fitting, we could estimate the bleed-through from red to green channel ranges from 3-8 ADU. It is negligible compared with EB1-eGFP pixel intensities in the range of 50-300ADU.

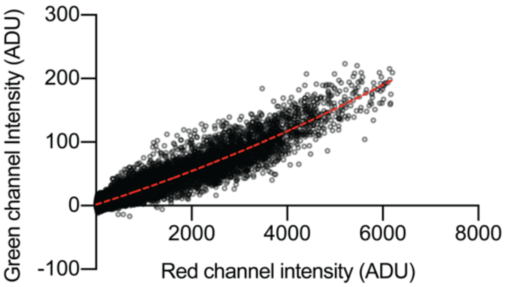


**Fig. S2** Bleed-through calibration curve. The measured intensity values are with camera offset removed. The red dashed curve indicates polynomial fitting with formula: $1.8+0.024x+1.3\times10^{-6}x^2$.

**<u>MC optimization method</u>**

The overall intensity profile of the EB1-eGFP comets within 0.5×4-8 μm$^2$ rectangular box can be fitted by $e^{-(y-\frac{\sigma^2/\lambda}{2})/\lambda}\,\mathrm{erfc}\left(-\frac{(y-\sigma^2/\lambda)}{\sqrt{2}\sigma}\right)$ obtained through convolution of gaussian with exponential decay, where $\sigma$ stands for the standard deviation of gaussian and $\lambda$ is the exponential decay length. The above fitted parameters $\sigma_{fit}$ and $\lambda_{fit}$ are used to set the range of $\sigma$ and $\lambda$ which are used as inputs for MC optimizations. We measured the mean $\sigma_{tetraspeck}$ and standard deviations $SD_{tetraspeck}$ of gaussian approximated point spread functions from tetra-speck beads as shown in Fig. S3. The range of input parameter $\sigma$ is determined by the intersection of $[\sigma_{fit} - 100nm, \sigma_{fit} + 100nm]$ and $[\sigma_{tetraspeck} - 2SD, \sigma_{tetraspeck} + 2SD]$. The range of input parameter $\lambda$ is determined by $[\lambda_{fit} - 60nm, \lambda_{fit} + 60nm]$.

To test the MC optimization method, we constructed model image generated by $I_{\mathrm{model}} = Ae^{-(y-\frac{\sigma^2/\lambda}{2})/\lambda}\,\mathrm{erfc}\left(-\frac{(y-\sigma^2/\lambda)}{\sqrt{2}\sigma}\right) + N$, where $A$ stands for the image intensity, $N$ stands for photon shot noise and background noise. The model images are created under three conditions: (A) $N =$

0, (B) max ($I_{\text{model}}$)~70 ADU and (C) max ($I_{\text{model}}$)~280 ADU. The output results $\sigma_{output}$ and $\lambda_{output}$ from MC optimization method as well as from analytical fitting are shown in Figs. S4 and S5. Both methods give similar predictions for $\lambda$ (Fig. Sx). MC optimization method performs better in predicting $\sigma$ compared with analytical fitting method as evidenced by the more compact cluster along *y*=*x* line shown in Fig. S4.

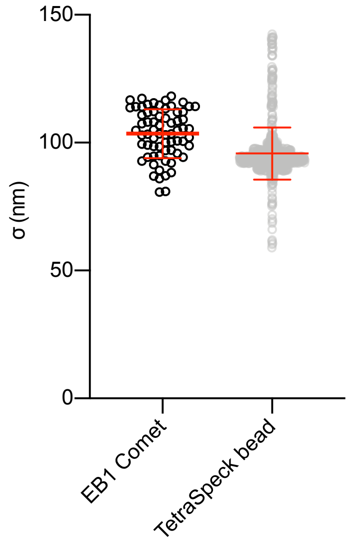


**Fig. S3** Experimentally measured σ of gaussian approximated point spread functions from EB1 comets profiles and tetraspeck beads. Mean and SDs for EB1 comets and tetraspeck beads are 104±10 nm (N=68) and 96±10 nm (N=806) respectively.

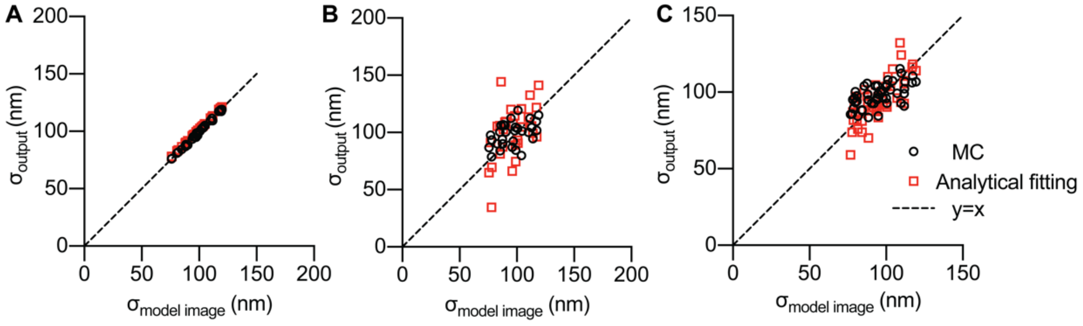


**Fig. S4** $\sigma_{output}$ for model images of (A) $N = 0$, (B) max ($I_{\text{model}}$)~70 ADU and (C) max ($I_{\text{model}}$)~280 ADU respectively. Output results from MC optimization are denoted by black circles and those from analytical fitting are denoted by red squares.

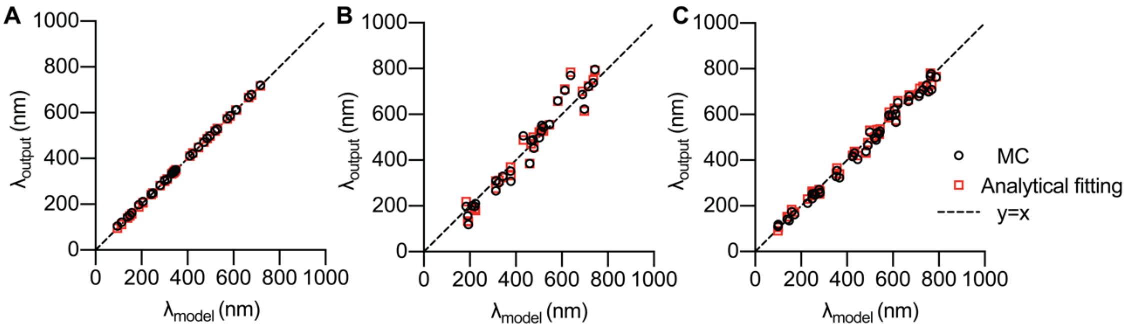


Fig. S5. $\lambda_{output}$ for model images of (A) $N = 0$, (B) max ($I_{\text{model}}$)~70 ADU and (C) max ($I_{\text{model}}$)~280 ADU respectively. Output results from MC optimization are denoted by black circles and those from analytical fitting are denoted by red squares.
